# Differences in the chitinolytic activity of mammalian chitinases on soluble and crystalline substrates

**DOI:** 10.1101/762336

**Authors:** Benjamin A. Barad, Lin Liu, Roberto Efrain Diaz, Ralp Basilio, Steven J. Van Dyken, Richard M. Locksley, James S. Fraser

## Abstract

Chitin is an abundant polysaccharide used by a large range of organisms for structural rigidity and water repulsion. As such, the insoluble crystalline structure of chitin poses significant challenges for enzymatic degradation. Vertebrates do not produce chitin, but do express chitin degrading enzymes. Acidic mammalian chitinase, the primary enzyme involved in the degradation of environmental chitin in mammalian lungs, is a processive glycosyl hydrolase that may be able to make multiple hydrolysis events for each binding event. Mutations to acidic mammalian chitinase have been associated with asthma, and genetic deletion of the enzyme in mice results in significantly increased morbidity and mortality with age. We initially set out to reverse this phenotype by engineering hyperactive acidic mammalian chitinase variants. Using a directed evolution screening approach using commercial fluorogenic substrates, we identified mutations with consistent increases in activity. To determine whether the activity increases observed with oligomeric substrates were consistent with more biologically relevant chitin substrates, we developed new assays to quantify chitinase activity with colloidal crystalline chitin, and identified a high throughput fluorogenic assay that gives sufficient signal to noise advantages to quantify changes to activity due to the addition or removal of a chitin binding domain to the enzyme. We show that the activity increasing mutations derived from our directed evolution screen were lost when crystalline substrates were used. In contrast, naturally occurring gain-of-function mutations gave similar results with oligomeric and crystalline substrates. We also show that the activity differences between acidic mammalian chitinase and chitotriosidase are reduced in the context of crystalline substrate, suggesting that previously reported activity differences with oligomeric substrates may have been largely driven by differential substrate specificity for the oligomers. These results highlight the need for assays against more physiological substrates when engineering complex metabolic enzymes, and provide a new approach that may be broadly applicable to engineering glycosyl hydrolases.

## Introduction

Polysaccharides are ubiquitous biopolymers that serve roles ranging from energy storage to signaling to structural rigidity (1–3). Polysaccharide catabolism is achieved by enzymes in the gylcosyl hydrolase, lytic polysaccharide monooxygenase, and glycosyltransferase families (4–6). Many polysaccharides assemble into higher order structures that complicate access by metabolic enzymes. Due to the high activation energy of effective binding to these structures, processive hydrolases are commonly employed by organisms to maximize the metabolism accomplished after each binding event (7, 8). In contrast to the complexity of polysaccharides in the environment, enzymatic hydrolysis is often quantified *in vitro* using small oligomer analogues, which release a chromophore or fluorophore when cleaved (9, 10). These simple substrates allow for high signal-to-noise and precise quantification, but activity measurement using these simple substrates is fundamentally limited in assessing enzyme activity on more complex substrates, due to their higher order, often crystalline, structures, variable polymer length, branching and other modifications. An additional challenge is quantifying processivity - processive hydrolases can cut bulk substrate multiple times for each binding event. As simple substrates often only have one site where hydrolysis will generate signal, often the degree of processivity cannot be assessed with these substrates, and the total activity of the enzyme with bulk substrate can be very different in the presence or absence of processivity (8).

The difficulties in assessing catabolism of complex polysaccharide substrates is exemplified by chitin, a ubiquitous polysaccharide comprised of ß-1,4-linked N-acetylglucosamine, that is produced by fungi and arthropods for structural rigidity and water repulsion (3, 11). Chitin polymers assemble into water-insoluble microcrystals, which have been observed in 3 different crystal forms, differentiated by the parallel or antiparallel orientation of neighboring chitin strands (12). Alpha-chitin, the most common and lowest energy conformation, forms antiparallel sheets that intercalate the N-acetyl groups of neighboring polymers and form tight hydrogen bonding networks (13). Strands of chitin must be extracted from this highly crystalline structure to be degraded, and the rate limiting step of catalysis has been observed to be the processive decrystallization of additional substrate from the bulk crystal (8, 14). This observation makes it particularly challenging to effectively associate degradation of short oligomeric analogues with true catalytic efficacy. Beyond this, the insolubility and recalcitrance of bulk chitin makes it a particularly challenging substrate to quantify with high precision. Recently, several new methods have tackled this problem by using labelled chitin substrates with chromatography (15, 16) as well as enzyme-coupled assays to generate colorimetric signal from reducing ends (17). These methods have enabled new insights into chitinase behavior, but their signal-to-noise ratio and throughput limit the ability to separate total activity into binding and catalysis, as well as other components of polysaccharide catabolism such as substrate specificity and processivity.

Dissecting chitinolytic activity to understand its mechanisms has particularly interesting implications for understanding how mammalian chitinases function in innate immunity. Chitin is not produced by mammals, but mammals have a conserved machinery to recognize and degrade environmental chitin that is inhaled or ingested (18, 19). The molecular mechanism of recognition of chitin and the signalling program it generates is not well understood, but breakdown of inhaled chitin is accomplished by the secreted enzymes acidic mammalian chitinase (AMCase) and chitotriosidase, which are conserved across mammals (18). Both are two domain family-18 glycosyl hydrolases consisting of a catalytic TIM-barrel domain and a C-terminal carbohydrate-binding domain. In AMCase, the two domains are connected by a 25 residue glycine- and serine-rich linker that is expected to be highly glycosylated, while chitotriosidase has a shorter, proline-rich linker that has also been found to be glycosylated (20–22). The roles of the linker and the C-terminal carbohydrate-binding domain in processing chitin have not been quantified.

AMCase is upregulated in response to chitin insult and is secreted into the airway lumen, where it interacts with crystalline chitin and breaks down the substrate (23). Consistent with the reported role of AMCase in asthma, there are polymorphisms of human AMCase (hAMCase) that increase its activity and have been associated previously with asthma protection (24). A trio of mutations found in humans, N45D, D47N, and R61M, which change residues to the wild type identities of mouse AMCase (mAMCase), has been previously described to increase specific activity against model substrates (25). Of these mutations, prior work has identified the R61M mutation as causing the largest increase in total activity, as well as the largest decrease in mice with the reverse M61R mutation (16). The mechanism by which these mutations alter binding and catalysis, both with simple and complex substrates, remains unclear. AMCase deficient mice accumulate chitin in their lungs and develop tissue fibrosis as an aging phenotype; external addition of recombinant chitinase to the airway reduces this phenotype (26). This suggests that AMCase is predominantly responsible for clearance of chitin from airways, and further suggests that enhancing AMCase activity may reduce chitin airway levels.

In this study, we initially tried to evolve variants of AMCase that would have enhanced activity to test the hypothesis that enhanced clearance of chitin would reduce the potential for age-related lung fibrosis. Our directed evolution approach was based on simple fluorogenic substrates. We found mutations that dramatically increase the activity of the enzyme by both improving binding and catalysis. We develop new approaches to quantifying bulk chitin degradation and discovered that these engineered mutations did not have the same effect with bulk substrates. We use these improved methods to assay the impact of the carbohydrate-binding domain on activity, and discover that it causes a minor K_m_ vs k_cat_ tradeoff but does not have a major effect on overall activity. We revert the asthma-protective mutants in the mouse background, and find that the dominant effect is a k_cat_ decrease from the M61R mutation. We also compare the activity of mAMCase and chitotriosidase with different small oligomeric substrates and with bulk chitin. These results highlight the need for assays against more physiological substrates when engineering complex metabolic enzymes, and provide a new approach that may be broadly applicable to engineering glycosyl hydrolases.

## Results

### Engineering of Hyperactive Chitinases

Recent efforts have identified recombinant chitinase as a potential direct therapy to ameliorate inflammatory lung symptoms that arise when native chitinase activity is compromised (26). To investigate whether we could improve the activity of mouse AMCase with the end goal of supplementing lung chitinase activity, we used error-prone PCR to generate libraries of mAMCase mutants (**Figure 1A**). Our recombinant expression approach, utilizing periplasmic secretion as described previously, also results in significant enzyme secreted into the media (27). Taking advantage of this, we assayed, in 96 well format, the ability of the spent media of individual mutants after protein expression to cleave 4MU-chitobiose. Comparing these results to both wild-type and engineered catalytically dead mutants, we found that while most mutations resulted in either total loss of protein activity or similar activity to wild-type, a small number of mutants were much more active than the wild-type (**Figure 1B)**. Because these assays were done directly on spent media, the activity for each well reports on the specific activity of the enzyme as well as to expression and secretion efficiency. To determine whether our results represented improvements in activity, we isolated and purified the two most active mutants: A239T/L364Q (**Figure 1D, pink**) was the most active mutant identified, with a 5-fold improvement in activity, and V246A (**Figure 1D, orange)**, which showed a 2-fold improvement in activity.

**Figure 1.**
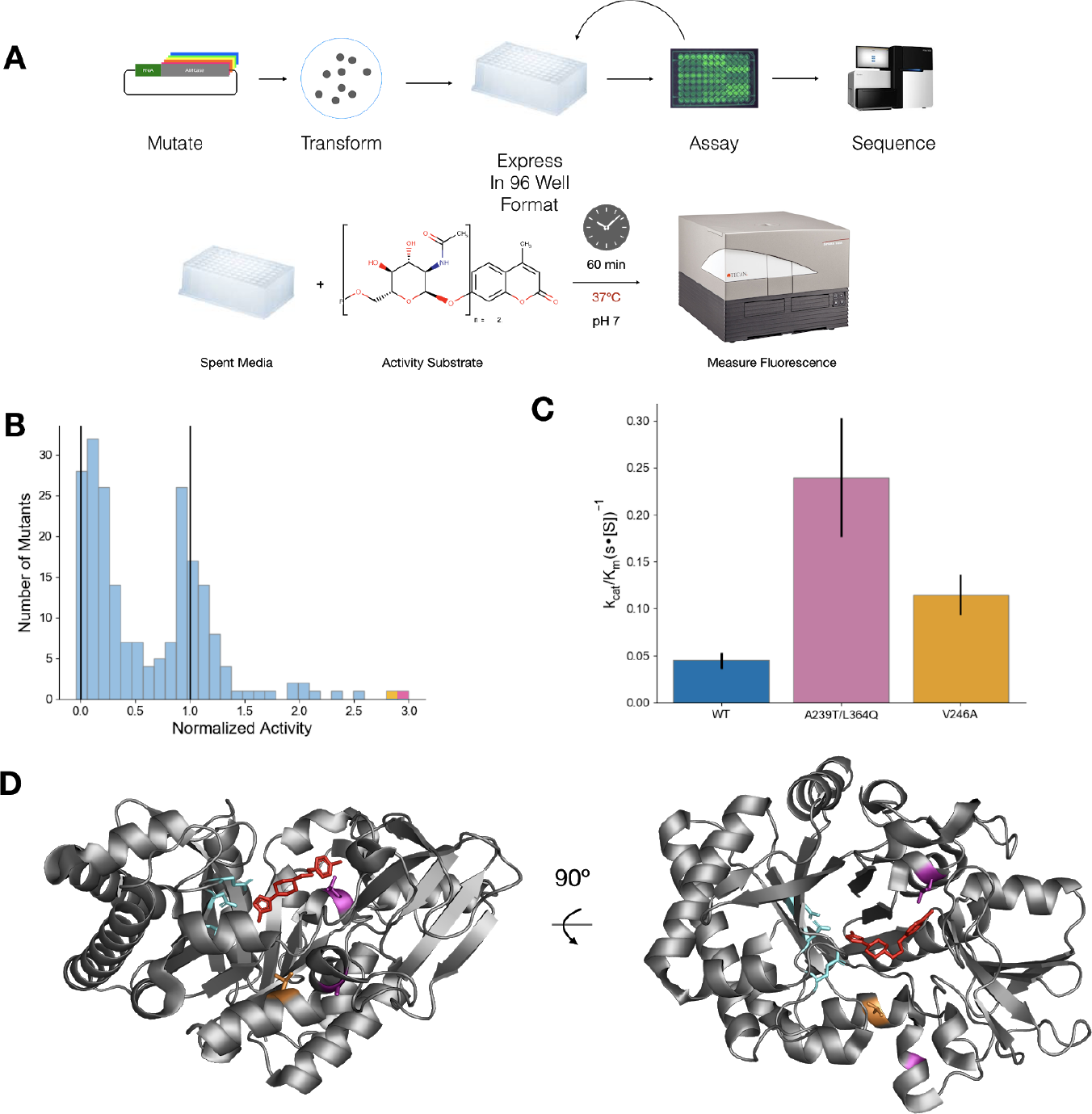
Engineering of hyperactive AMCase mutants. (a) Workflow for directed evolution of AMCase. Mutants of AMCase were generated via error-prone PCR, then transformed and grown out from individual colonies in 96-well blocks. After expression, activity was measured using the 4MU-chitobiose substrate incubated with the expression media. (b) Distribution of activity for mutants with 1-3 mutations per construct. Vertical lines at 0 and 1 represent a catalytically dead negative control and a wild type positive control, respectively. The best two results are highlighted in purple and orange. (c) k_cat_/K_m_ of purified hyperactive mutants using the 4MU-chitobiose assay (d) Structure of AMCase catalytic domain highlighting A239T/L364Q (orange) and V246A (blue). The active site catalytic network is highlighted in teal, and an inhibitor that binds to the active site cleft is shown in red.

After purification, we measured the specific activity of the assay using a one-pot continuous-read fluorescent assay based on the previously developed enzyme-coupled assay (17), and replicated the improvements observed in the unpurified screening format (**Figure 1C**). Both mutants improved significantly in k_cat_, while the A239T/L364Q had a nonsignificant improvement in K_m_ (**Table 1**). These results can be rationalized by the locations of the mutations in the structure. A239T and V246A are relatively distant from the active site, but L364Q is positioned immediately at the binding site for chitin (**Figure 1D**).

**Table 1.**
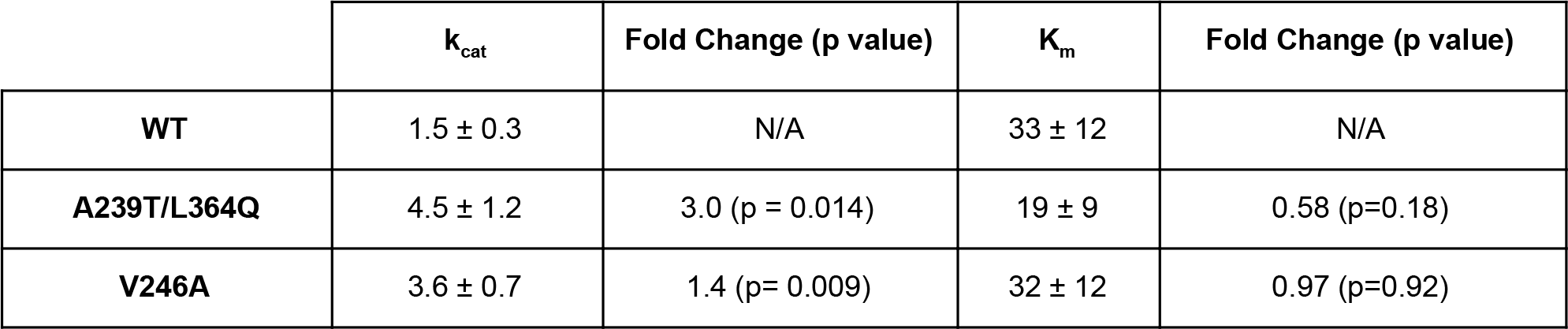
rates for engineered mutants. k_cat_ values are reported in units of 1/s. K_m_ values are reported in units of mM for 4MU-chitobiose and % w/v for chitO assays. Fold changes are relative to wild type enzyme.

| | $k_{\text{cat}}$ | Fold Change (p value) | $K_m$ | Fold Change (p value) |
| --- | --- | --- | --- | --- |
| **WT** | $1.5 \pm 0.3$ | N/A | $33 \pm 12$ | N/A |
| **A239T/L364Q** | $4.5 \pm 1.2$ | 3.0 (p = 0.014) | $19 \pm 9$ | 0.58 (p=0.18) |
| **V246A** | $3.6 \pm 0.7$ | 1.4 (p= 0.009) | $32 \pm 12$ | 0.97 (p=0.92) |

### Comparison of the activity of the catalytic domain of AMCase to the full length enzyme with new approaches

An alternative hypothesis for the increased activity is that the engineered variants have high specificity for the fluorophore or a smaller oligomer. This idea motivated us to develop new assays on larger and more complex crystalline chitin material. As a first control, we wanted to assess the contribution of the catalytic and carbohydrate-binding domain of AMCase. Due to the short length of the oligomeric substrate, a reaction with the small 4MU substrate is likely to be driven only by local interactions in the catalytic domain and the presence of the carbohydrate-binding domain should not affect the reaction rate. In contrast, the carbohydrate-binding domain has been hypothesized to play a role in binding crystalline chitin (28, 29).

We expressed and purified the isolated catalytic domain of AMCase, as well as the full length enzyme, using an *E. coli* periplasmic expression approach (27). We first measured the ability of the enzyme to catalyze the breakdown of 4-methylumbelliferone (4MU) conjugated chitobiose, using a continuous read approach at pH 7. The activities of the two constructs were indistinguishable, either in binding or catalysis (**Figure 2A**, **Table 2**, p=0.3).

**Figure 2.**
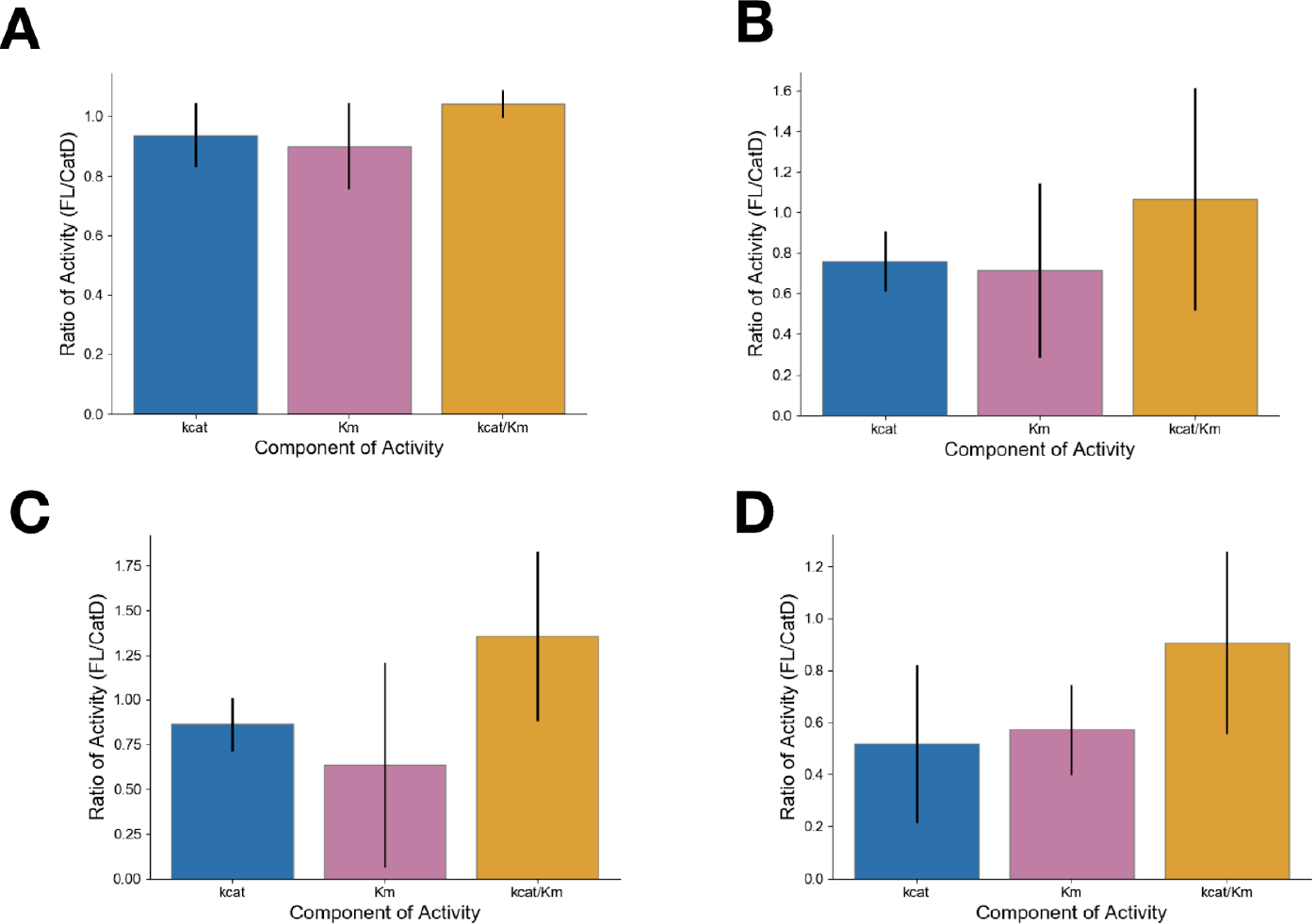
Activity comparisons of AMCase catalytic domain and full length enzyme. Difference in k_cat_,K_m_, and k_cat_/K_m_ of AMCase catalytic domain and full length enzyme generated via (a) 4MU-chitobiose assay (b) colloidal chitin clearance assay (c) reducing sugar generation assay quantified with potassium ferricyanide (d) chitooligosaccharide oxidase coupled peroxidase assay. Error bars denote propagated standard deviation of fit (accounting for covariance).

**Table 2.** Calculated rate constants of AMCase catalytic domain and full length enzyme k_cat_ values are reported in units of 1/s. K_m_ values are reported in units of mM for 4MU assays and % w/v for colloidal clearance, ferricyanide, and chitO assays.

|  | Catalytic Domain |  | Full Length Enzyme |  |
| --- | --- | --- | --- | --- |
| | $k_{cat}$ | $K_m$ | $k_{cat}$ | $K_m$ |
| **4MU-Chitobiose** | $1.12 \pm 0.09$ | $28 \pm 3$ | $1.05 \pm 0.07$ | $25 \pm 2$ |
| **Colloidal Clearance** | $0.00140 \pm 0.00008$ | $0.09 \pm 0.03$ | $0.00106 \pm 0.00002$ | $0.07 \pm 0.02$ |
| **Ferricyanide** | $0.454 \pm 0.042$ | $0.046 \pm 0.02$ | $0.39 \pm 0.02$ | $0.029 \pm 0.007$ |
| **ChitO** | $0.9 \pm 0.1$ | $0.033 \pm 0.006$ | $0.54 \pm 0.08$ | $0.017 \pm 0.005$ |

To quantify the role of the carbohydrate-binding domain, we developed a new methodology for quantifying chitinase activity with complex substrates. We took advantage of the commercial availability of colloidal chitin substrates, which are more uniform in size and shape and to have reduced settling times compared to other substrates shrimp shell chitin. We first attempted to measure colloidal chitin hydrolysis by the disappearance of scattering by solid substrate as it is converted into small oligomeric products. We could not distinguish a statistically significant difference between the two variants with this approach, which was likely limited by the relatively small dynamic range and large amount of enzyme required to produce a measurable change in scattering (**Figure 2B**, **Table 2**, p=0.8). Each cutting event only minimally alters the scattering of chitin crystals, and many cuts are likely necessary to fully solubilize crystals.

We next attempted to quantify the production of soluble reducing ends, which we hypothesized would more more sensitively report individual catalytic events, resulting in an overall improvement in signal. The first method we used to assay production of soluble reducing ends was a ferricyanide reduction assay (30): after incubating colloidal chitin with AMCase at 37ºC for up to 18 hours, we quenched the reaction with sodium carbonate, removed the insoluble chitin by centrifugation, and quantified the non-enzymatic reaction of soluble reducing sugars with potassium ferricyanide, read out by the disappearance of the yellow color by absorbance at 420 nm. With this assay, we were not able to identify a significant difference in total activity, but were able to identify that the inclusion of the carbohydrate-binding domain created a small improvement in K_m_ that was offset by a reduction in the k_cat_ of AMCase (**Figure 2C**, **Table 2**, p=0.2). This tradeoff did not result in a large difference in activity. Moreover, the endpoint-based requirements of the assay and of the dynamic range available in measuring reduction in absorbance were limiting. We next developed a new assay based on previous work using chitooligosaccharide oxidase (chitO) in combination with horseradish peroxidase to generate signal specifically from the production of chitin reducing ends (17). To convert this assay from endpoint to continuous readout, we took advantage of fluorogenic substrates for horseradish peroxidase and carefully washed the colloidal chitin to enable signal measurement without removal of the insoluble component. This gain-of-signal fluorescent assay had much improved signal-to-noise and sensitivity, and improved quantification of the kinetic parameters of chitinase activity. Using this assay, we were able to more confidently determine the tradeoff between improved binding (p=0.02) and loss of maximal catalytic activity (p=0.009) with the inclusion of the carbohydrate-binding domain, which resulted in no significant change in total activity (**Figure 2D**, **Table 2**, p=0.8).

### Small substrates can be misleading for engineered chitinases

Having this new assay in hand, we tested whether the activity increases observed with the 4MU-chitobiose mutant resulted in similar improvements to degradation of bulk chitin. Using purified protein, we measured the activity of the mutants to degrade colloidal chitin using the enzyme-coupled chitO assay, and discovered that the A239T/L364Q mutant had lost all measurable activity, while the V246A mutant was not statistically significantly more active than the wild type (**Figure 3**). The loss of activity of the double mutant suggests that the improvements were driven by the L364Q mutation interacting with the 4MU fluorophore, which can be rationalized structurally (**Figure 1D**). The stark difference in results between the results with the 4MU and chitO assay underscores the need for assays of catabolism of bulk chitin substrate, even during the initial stages of screening.

**Figure 3.**
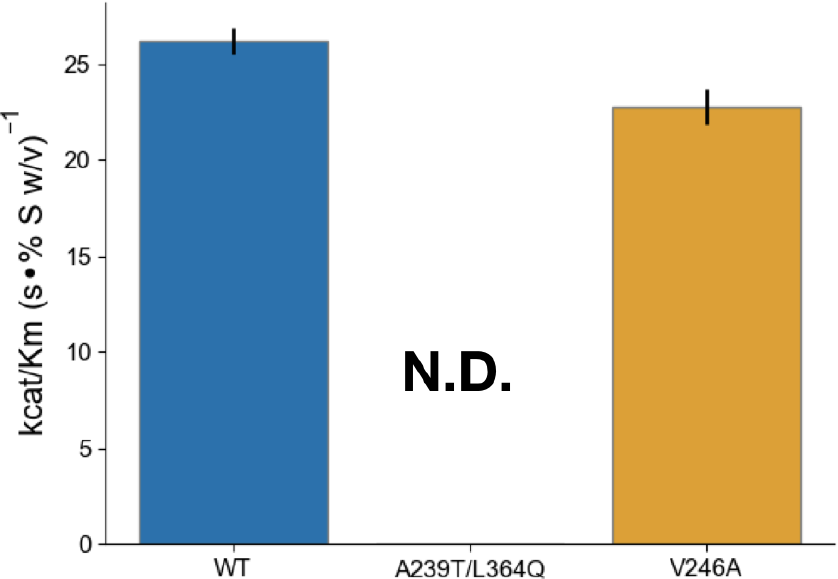
Engineered mutant activity with the novel chitooligosaccharide oxidase assay. Difference in k_cat_/K_m_ of purified hyperactive mutants using the 4MU-chitobiose assay. k_cat_ values are reported in units of 1/s. K_m_ values are reported in units % w/v. Error bars denote propagated standard deviation of fit (accounting for covariance). The A239T/L364Q mutant had too little total activity to measure k_cat_ or K_m_.

### Effects of Human Asthma-Associated Mutants in the Mouse context

Motivated by the result on the engineered mutations, we wanted to test naturally occurring mutations that have previously been shown to have different activities using the 4MU assay. We focused on a trio of mutations in AMCase in humans, N45D, D47N, and R61M, that confer significantly increased activity to AMCase (16, 25). In all three cases, the identity of the mutated residues becomes the same as the identity of the residues of the mouse wild-type protein. To better understand the mutational landscape between the mouse and human enzymes, which have 81% sequence identity and differ by 92 total polymorphisms, we made the reverse mutations in the mouse background to quantify their effect on activity using both 4MU-chitobiose and bulk chitin. First we measured the activity of the mutations using 4MU-chitobiose, which showed that the mouse wild-type residues were more active than the human wild-type residues, similar to previous results in the human background. The activity difference between wild-type and the M61R mutant was caused by a decrease in k_cat_ and a small increase in K_m_ (**Figure 4A**, **Table 4**, p=0.03). Smaller effects were observed for the individual D45N and N47D mutations, but the effects were reversed by the charge-swapped D45N/N47D construct. The full triple mutant was the least active (p=0.001). These results suggest that the different residue identities have overall very similar effects in the mouse and human backgrounds, and that the low-activity wild-type identities in primates may have arisen due to reduced selective pressure for chitinase activity around pH 7. To understand to what degree these effects observed with the oligomeric substrate are relevant to enzyme activity on bulk chitin, we assayed the activity of humanizing mutations in mAMCase using the enzyme-coupled chitO assay. The results were similar to those using the 4MU substrate, with the largest effect of any individual mutation and the majority of the effect of the triple mutation contributed by the M61R mutant (**Figure 4B**, **Table 4**, p=0.02). The effects of the D45N and N47D mutations were less pronounced in the chitO background, while the M61R mutation had a similar effect on both K_m_ and k_cat_.

**Figure 4.**
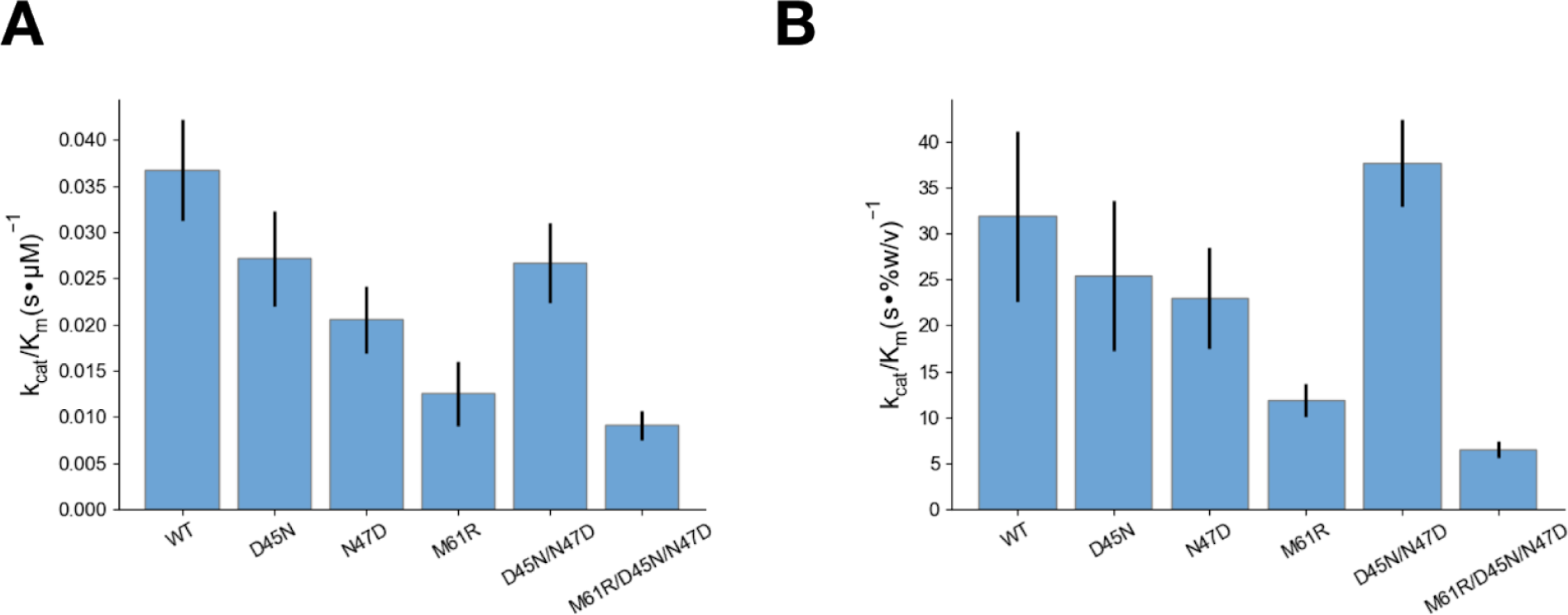
Comparison of activity of AMCase asthma-associated mutants. Measurement of k_cat_/K_m_ for reversed asthma-associated mutants in the mouse background using the (a) 4MU-chitobiose and (b) chitO assays. Error bars denote propagated standard deviation of fit (accounting for covariance).

**Table 3.** Calculated rate constants of engineered mutants with k_cat_ values are reported in units of 1/s. K_m_ values are reported in units % w/v. Fold changes are relative to wild type enzyme.

| | $k_{cat}$ | Fold Change (p value) | $K_m$ | Fold Change (p value) |
| --- | --- | --- | --- | --- |
| **WT** | $0.78 \pm 0.05$ | N/A | $0.030 \pm 0.002$ | N/A |
| **A239T/L364Q** | N.D. | N/A | N.D. | N/A |
| **V246A** | $0.80 \pm 0.07$ | 1.03 (p=0.7) | $0.035 \pm 0.004$ | 1.17 (p=0.1) |

**Table 4.** Calculated rate constants for AMCase and chitotriosidase k_cat_ values are reported in units of 1/s. K_m_ values are reported in units of mM for 4MU-chitobiose and % w/v for chitO assays.

|  | 4MU-Chitobiose |  | chitO |  |
| --- | --- | --- | --- | --- |
| | $k_{cat}$ | $K_m$ | $k_{cat}$ | $K_m$ |
| **WT** | $1.1 \pm 0.1$ | $30 \pm 3$ | $1.0 \pm 0.2$ | $0.032 \pm 0.008$ |
| **D45N** | $0.94 \pm 0.07$ | $35 \pm 6$ | $0.8 \pm 0.1$ | $0.032 \pm 0.008$ |
| **N47D** | $0.71 \pm 0.05$ | $35 \pm 6$ | $0.74 \pm 0.03$ | $0.032 \pm 0.008$ |
| **M61R** | $0.8 \pm 0.2$ | $60 \pm 12$ | $0.48 \pm 0.05$ | $0.041 \pm 0.005$ |
| **D45N/N47D** | $0.98 \pm 0.09$ | $37 \pm 5$ | $1.02 \pm 0.04$ | $0.027 \pm 0.003$ |
| **D45N/N47D/**<br>**M61R** | $0.46 \pm 0.05$ | $50 \pm 7$ | $0.31 \pm 0.04$ | $0.047 \pm 0.006$ |

### Comparison of Acidic Mammalian Chitinase and Chitotriosidase

Next we wanted to compare AMCase to the other major human chitinase, Chitotriosidase. Both enzymes are expressed in lungs, but only acidic mammalian chitinase is strongly overexpressed in response to chitin insult (31). Previous reports using the 4MU assay have indicated activity differences and no synergistic effects (32), but this result is convolved with the substrate specificity of dimer and trimer chitin oligomers. Whether AMCase and Chitotriosidase have similar activity on crystalline substrates has not been previously examined.

We sought to understand how differences in binding, substrate specificity, and hydrolytic activity differed between the two enzymes. We investigated substrate specificity by comparing the ability of each enzyme to cleave the terminal glycosidic linkage on 4MU-chitobiose and 4MU-chitotriose, representing hydrolysis in different substrate binding poses to generate chitobiose vs chitotriose as a substrate. When assayed the 4MU-chitobiose substrate, AMCase had more than double the activity of chitotriosidase, driven by a significant difference in K_m_ (**Figure 5A**, **Table 5**, p=0.01). In contrast, the 4MU-chitotriose substrate led to tighter binding for both AMCase and chitotriosidase, but the difference was much larger with chitotriosidase, leading to a smaller gap in activity between the two enzymes (**Figure 5B**, **Table 5**). The reduction in observed k_cat_ for both enzymes was likely driven by the alternative, non-fluorogenic reaction trajectory in which the 4MU-chitotriose is cleaved into chitobiose and 4MU-bound N-acetylglucosamine, leading to a systematic underestimate of k_cat_. The differences in the K_m_ suggests that chitotriosidase benefits more from the extended binding interactions available with the larger 4MU-chitotriose substrate. We next assayed the differences in activity with a bulk substrate using the chitO coupled assay. The difference in activity was much smaller in this assay, with the majority of the activity difference being driven by k_cat_ differences (**Figure 5C**, **Table 5**, p=0.006). These results further highlight the differences in substrate-specific effects between chitotriosidase and acidic mammalian chitinase as had been previously described (32), and suggest that much of this difference can be attributed to differential binding efficiency for short chitin oligomers.

**Figure 5.**
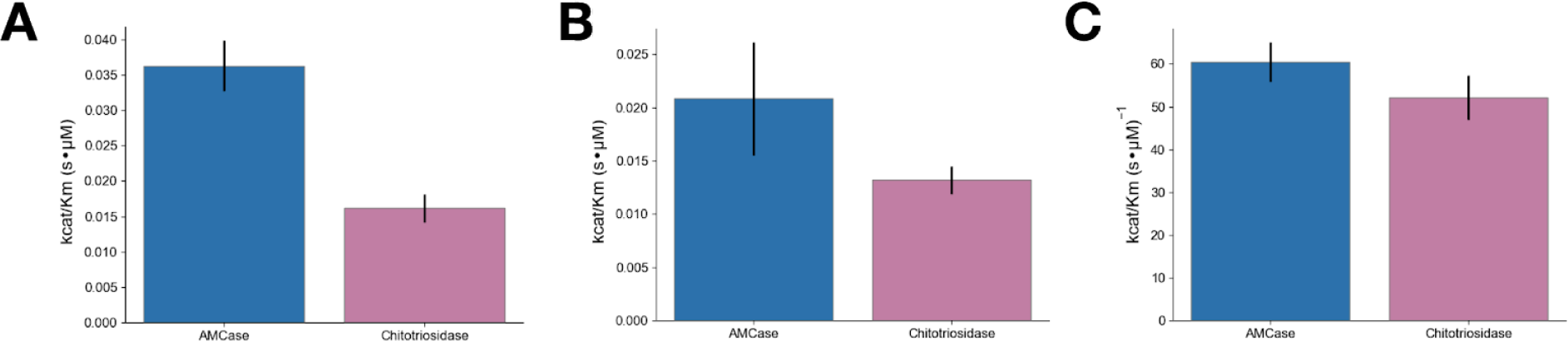
Comparison of AMCase and Chitotriosidase. Differences in k_cat_/K_m_ between AMCase (green) and Chitotriosidase (orange) using (a) 4MU-chitobiose (b) 4MU-chitotriose (c) chitooligosaccharide oxidase coupled peroxidase assay. Error bars denote propagated standard deviation of fit (accounting for covariance).

**Table 5.** Calculated rate constants for AMCase and Chitotriosidase k_cat_ values are reported in units of 1/s. K_m_ values are reported in units of mM for 4MU assays and % w/v for chitO assays.

|  | AMCase |  | Chitotriosidase |  |
| --- | --- | --- | --- | --- |
| | $k_{cat}$ | $K_m$ | $k_{cat}$ | $K_m$ |
| **4MU-chitobiose** | $1.02 \pm 0.05$ | $28 \pm 4$ | $1.1 \pm 0.1$ | $68 \pm 15$ |
| **4MU-chitotriose** | $0.5 \pm 0.1$ | $24 \pm 11$ | $0.33 \pm 0.03$ | $25 \pm 4$ |
| **ChitO** | $1.06 \pm 0.06$ | $0.018 \pm 0.002$ | $0.82 \pm 0.05$ | $0.016 \pm 0.003$ |

## Discussion

Broadly, these results demonstrate the value of quantifying chitinase kinetics with bulk substrates with the same care used with model substrates (fluorogenic oligomers). Our results suggest that the effectiveness and sensitivity of the one-pot chitooligosaccharide oxidase coupled assay makes it an ideal approach for monitoring chitinase activity. While in some cases the results of the activity assays closely resembled the 4MU-chitobiose assays, in others the activities were tremendously different, underscoring the need for quantitative measures of bulk chitin catabolism. This proved to be particularly true for studies of the effects of multiple domains, which necessarily cannot bind the same short oligomer the same way they could a chitin crystal, as well as for engineered variants, in which screening with short fluorogenic substrates led to artifacts that may be related to fluorophore binding. The sensitivity and throughput available with the chitO coupled assay enables more precise and quantitative measurements of bulk chitin catabolism than was previously available, and we expect that this technique will be effective for deconstructing different aspects of enzyme activity.

In contrast to the majority of cases, which had reasonable agreement between the bulk experiments and the small oligomers, our efforts to engineer hyperactive chitinases were limited by the use of the 4MU-chitobiose substrate as a screening tool. Our best mutants from screening had significant increases in activity, but once the purified mutants were assayed by the chitO assay, the improvements were not present. In the case of the A239T/L364Q mutant, there was no quantifiable activity with bulk substrate. The classic maxim is that “in protein engineering you get what you screen for”, and in this case that was maximizing binding efficiency for the 4-methylumbelliferone fluorophore, as well as the smaller substrate overall. The result underscores the need in the future for utilizing frequent counter-screening with bulk chitin when performing selection experiments for chitin processing. One challenge to accomplishing this is that, while the chitO assay is more sensitive and high throughput than previous techniques, it is sensitive to free sugars and other components of the media that limits its utility for direct screening. It is possible that with small scale purification using nickel resin, we may in the future be able to directly screen activity of mutants using the chitO method. In combination with recent advances in guiding small library directed evolution with machine learning (33), we may be able to effectively use this approach to find hyperactivating mutants without the requirement of using chitobiose substrates.

Quantitatively, with the exception of the engineered mutants, the kinetic parameters measured with the 4MU and bulk chitin assays were well aligned, with k_cat_ values that were remarkably similar, suggesting that the 4MU assay effectively captures the chemical step of hydrolysis, and K_m_ values that were on the order of 30 μM for the 4MU-chitobiose and 0.03% w/v for the bulk chitin assay. Under the approximation of infinite polymer length, there is one potential binding site per monomer of N-acetylglucosamine. Each chitin monomer unit has a molecular mass of 203.21 g/mol, so 0.03% w/v or 0.3 g/L would correspond to approximately 1.5 mM, 50 times greater than the K_m_ observed for the small oligomeric substrates. We hypothesize that the higher effective K_m_ reports on the relative crystallinity of the chitin, with a small proportion of theoretical substrate binding sites being accessible to the enzyme. In the future, it may be possible to alter this crystallinity, using partial deacetylation, co-application of chitin-binding enzymes that might loosen the crystalline geometry, or physical milling to alter the surface area to volume ratio.

Using the new bulk activity measurements, we were able to discern a tradeoff between k_cat_ and K_m_ with the addition of the carbohydrate-binding domain of AMCase, as K_m_ improved from 0.0333% ± 0.0056% to 0.0172%± 0.0049% chitin w/v (p=0.02), while k_cat_ decreased from 0.944 ± 0.111 1/s to 0.540 ± 0.083 1/s (p=0.007). While the improved binding with the addition of additional binding sites for chitin is unsurprising, the difference in the k_cat_ is less clear. Previous work has suggested that, for some chitinases, the rate limiting step in bulk catalysis is processivity (14). This result appears to support that hypothesis for AMCase as well, since the additional binding motif may inhibit the ability of the catalytic domain to effectively slide to new binding sites. If AMCase processivity proves to be rate limiting, given the closely matched k_cat_ for 4MU-chitobiose, in which processivity is not possible and cannot be rate limiting, and bulk chitin, it suggests that the rates of catalysis and processivity may be very similar in the mouse enzyme. This may be a result of selection optimizing the overall rate of the enzyme or the relative size of products generated by the enzyme. For example, larger oligomers could be produced if decrystallization and sliding were much faster than the rate of hydrolysis. Additionally, these larger oligomers may be the relevant molecules sensed by the mammalian immune system, as seen in plants (34). The carbohydrate-binding domain may further impact other aspects of catalysis, such as selecting specific chitin local morphology, binding chitin in the correct orientation, modulating processivity, or releasing when strands of chitin become too short to further process. The secondary domain may also impact the stability of folding of the catalytic domain, leading to incorrect assessment of concentration of folded protein and deflation of measured k_cat_. Additionally, the assayed constructs lack post-translational modifications. Acidic mammalian chitinase is predicted to have multiple O-linked glycosylation sites in the linker between the catalytic domain and the carbohydrate-binding domain (22), which may have significant effects on interactions with crystalline substrates.

The methods developed here can give information about binding and catalysis with relevant substrates, but questions still remain about processivity, endo vs exo preference, and potential clustering and cooperative behavior between multiple enzymes. One avenue to more fully characterize these aspects of catalysis will be single-molecule measurements of kinetics. Recently, significant progress has been made in measuring chitinase activities by single-molecule microscopy (14, 35, 36), and applying this approach to mammalian chitinases, ideally with native glycosylation, may help to break down the effects of different mutations on activity, give new insights into the function of the carbohydrate-binding domain, and help to differentiate the enzymatic role of chitotriosidase and acidic mammalian chitinase.

## Experimental Procedures

### Protein Preparation

Constructs expressing a fusion of a protein A secretion sequence targeting periplasmic expression, AMCase or chitotriosidase, and a C-terminal V5-6xHIS as previously described (27) were ordered from Atum. Mutants of AMCase were generated via PCR mutagenesis. Chitinase-containing plasmids were transformed into BL21 cells and expressed overnight in ZY Autoinduction media at 37°C for 3 hours followed by 19°C overnight. We added protease inhibitor at the temperature change to minimize proteolysis of periplasmically expressed protein. Pelleted cells were lysed via osmotic shock in a two step procedure. First, cells were resuspended in 20% Sucrose w/v, 20 mM Tris pH 6.5, 1mg/mL lysozyme, 1 µL universal nuclease, with a protease inhibitor tablet. The resuspended cells were incubated at 37°C for 1 hour, then pelleted via centrifugation at 15000 RCF for 15 minutes. The supernatant was collected, and the pellet was resuspended in a wash buffer of 20 mM Tris pH 6.5 and 150 mM NaCl and incubated for 15 minutes at 4°C. The cells were centrifuged at 15000 RCF for 15 minutes and the supernatant was combined with the supernatant from the first step to form the combined lysate. The combined lysate was bound to a HisTrap FF column, washed with 100 mM Tris pH 6.5, 150 mM NaCl, then eluted with a gradient into 100 mM Tris pH 6.5, 150 mM NaCl, 500 mM imidazole. Fractions were selected for further purification based on activity assay with a commercial fluorogenic substrate (described below). Active fractions were pooled and subject to dialysis overnight into 100 mM Sodium Acetate pH 4.5, 150 mM NaCl, 5% glycerol w/v followed by filtration to remove insoluble aggregate and dialysis into 100 mM Tris pH 6.5, 150 mM NaCl, 5% glycerol w/v. The protein solution was concentrated and separated via size exclusion chromatography on a Superdex S75 16/600. Fractions were selected based on purity as assessed via SDS-page gel electrophoresis, and based on activity as assayed with a commercial fluorogenic substrate.

### Analysis of Kinetic Data

Kinetic measurements were made in a range of substrate concentrations outside of pseudo-first-order conditions. To robustly measure rates of catalysis, we fit our data using non-linear least-squares curve fitting to simple relaxation models for enzyme kinetics:

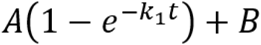

Where A shows the asymptotic signal from the clearance of substrate, k_1_ is the rate constant of relaxation, and B is the background signal of the assay condition. To this end, we developed a small free and open source python library for relaxation modeling, which is available on GitHub: https://github.com/fraser-lab/relax. Generally a single-step relaxation model was required, but in cases where residuals showed significant structure, additional steps were added as either relaxation or linear fits (in cases where kinetics were pseudo-first-order). The product of A and k_1_ yields a rate appropriate for k_cat_/K_m_ determination. Specific data analysis scripts using relax.py are available at https://github.com/fraser-lab/chitin_analysis.

### Continuous fluorescence measurements to quantify activity using commercial oligomeric substrates

Catalytic activity was assayed using 4-methylumbelliferyl chitobiose and 4-methylumbelliferyl chitotriose as described previously (37) with one critical modification. 10 nM chitinase enzyme was incubated with varying concentrations of 4MU-chitobiose or 4MU-chitotriose up to 433 μM in McIlvaine Buffer(38) pH 7 at 37° C. The 4-methylumbelliferone (4MU) fluorophore is quenched by a ß-glycosidic linkage to a short chitin oligomer, which is cleaved by a chitinase enzyme, which generates fluorescence with peak excitation at 360 nm and emission at 450 nm. Previously, the reaction was quenched and the pH was raised to maximize the quantum yield of the 4MU substrate. To avoid noise introduced by quenching and substrate concentration, we instead measured fluorescence at regular intervals during the course of the reaction without a pH shift and determined the rate using a single step relaxation model. This allowed us to measure rates of catalysis under a large range of conditions without needing to account for the proper time to quench to maximize signal without the reaction reaching completion. The processing for data collected from this assay is illustrated in **Figure S1**.

### Bulk Clearance Activity Assay

Borohydride-reduced colloidal chitin was purchased as a powder from Megazyme and resuspended to 4% w/v in pH 7 McIlvaine buffer. Higher concentrations did not stay in suspension effectively. To remove soluble chitin oligomers, the suspension was pelleted by centrifugation at 3200 RCF, the supernatant was discarded, and the pellet was resuspended in McIlvaine buffer. This wash step was repeated a total of 5 times. A concentration series was prepared by serial dilution of this washed 4% w/v stock in McIlvaine buffer, and 50 µL of each substrate concentration was incubated with 50 µL of 200 nM chitinase at 37° C in a clear-bottomed 96-well microplate with a lid that was sealed around the sides with parafilm to minimize evaporation. Clearance of substrate was monitored by reduction of scattering at OD_680_ for 72 hours with shaking between reads to maintain substrate suspension. The processing for data collected from this assay is illustrated in **Figure S2**.

### Potassium Ferricyanide Reduction Assay

4% w/v colloidal chitin was washed as above, then diluted serially to generate a concentration range from was incubated with 1-100 nM chitinase for up to 18 hours at 37°C. At the endpoint of incubation, 50 μL of reaction mixture was quenched by the addition of 100 μL of 400 mM sodium carbonate. The insoluble chitin was pelleted by centrifugation at 4000 RCF, then 100 μL of supernatant was mixed with 100 μL of 0.6 g/ml potassium ferricyanide in a 96 well microplate with clear bottoms and a lid that was sealed around the sides with parafilm to minimize evaporation. The microplate was incubated for 4 hours at 42°C to maximize the rate of the non-enzymatic reduction of potassium ferricyanide by solubilized reducing sugars. During incubation absorbance at 420 nm was read out in 1 minute intervals. We found that progress curve analysis gave results for this data, and instead ultimately found the difference between the maximum and minimum absorbance to be a more robust measure of total reducing sugar generation in the 18 hour incubation with chitinase. The processing of the data for this assay to generate rates is illustrated in **Figure S3**.

### Chitooligosaccharide Oxidase Coupled Peroxidase Assay

Processing of colloidal chitin and resultant generation of new reducing sugar moieties was monitored, as previously described,(17) by oxidation by chitooligosaccharide oxidase (ChitO), producing as a byproduct peroxide, which in turn is converted into a fluorescent signal by horseradish peroxidase (HRP) and QuantaRed peroxidase substrate.(39) ChitO was purchased from Gecco Biotech, HRP and QuantaRed substrate were purchased from Sigma. Incorporating a fluorogenic HRP substrate improves the dynamic range of the experiment and enables real time observation of reducing sugar cleavage in a one-pot reaction incorporating insoluble chitin, chitinase, chitO, HRP, and QuantaRed substrate. Briefly, a 50 μL solution containing 1-10 nM chitinase, 20 U/mL HRP, 100 nM ChitO, 0.5 μL of QuantaRed substrate, and 10 μL of QuantaRed enhancer solution in McIlvaine buffer pH 7 was mixed with 50 μL of washed colloidal chitin substrate, as prepared above, in a black 96-well microplate with a lid to minimize evaporation. The plate was incubated with at 37° C and the fluorescence of the QuantaRed substrate was measured at 1 minute intervals for 16 hours. The progression of fluorescence over time was modeled as a relaxation process as described above, after subtracting the signal from a chitinase-free control, which had significant signal that was modulated by the washing of the colloidal chitin. This enzyme-coupled reaction proved to be very sensitive to reaction conditions, with artifacts introduced by insufficient excess of chitO or HRP as well as by insufficiently washed colloidal chitin. With careful washing of the colloidal chitin and sufficient prewarming of both enzyme and substrate solutions, rates can be reliably measured for chitin concentrations ranging from 0.0005% to 2% colloidal chitin w/v, and for chitinase concentrations as low as 50 pM. The processing of data from this experiment is illustrated in **Figure S4**.

### Random Mutagenesis and Screening

Random mutations were generated using the commercial Genemorph II random mutagenesis kit.(40) Briefly, the catalytic domain of acidic mammalian chitinase was amplified via error-prone PCR with varying amounts of parent plasmid present. We titrated the amount of parent plasmid until each clone carried 1-2 mutations. We then performed restriction digestion using StyI and Eco130I and ligation using Quick Ligase to generate plasmids containing our mutations. We transformed these into electrocompetent BL21(DE3) *E. coli*. Individual colonies were picked and grown overnight in 96-well deep-well blocks, then 20 μL of starter media was used to inoculate 300 μL of ZY media in deep well blocks, which was then used to express the protein at 30°C overnight. After expression, 50 μL of media from individual wells was mixed with 50 μL of 21.6 μM 4MU-chitobiose in McIlvaine buffer pH 7, which had been prewarmed to 37°C. The mixture was monitored by fluorescence as described above, and compared to positive and negative controls which had been expressed in the same plate. Mutants with increased activity were grown out, mini-prepped, sequenced, retransformed, and expressed and rescreened in this manner in triplicate to confirm improved activity. Winners at this point were stored individually and pooled for further error-prone PCR and screening.

---

**Figure S1.**
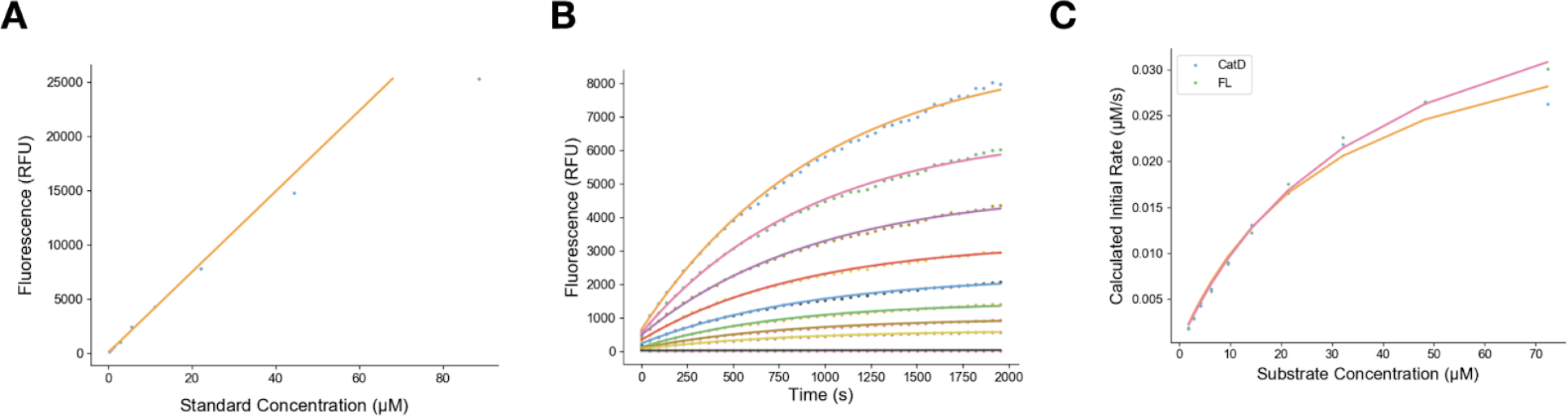
Processing 4MU assay data (a) Standards of 4MU were measured by fluorescence at 360 nm excitation and 420 nm emission and concentrations below 50 μM fit well to a linear regression (b) Progress curves of a concentration series of 4MU-chitobiose were fit by a non-linear relaxation analysis to extract initial rates. (c) Initial rates were plotted against substrate concentration and were fitted via non-linear regression to a Michaelis-Menten curve to extract rate constants.

**Figure S2.**
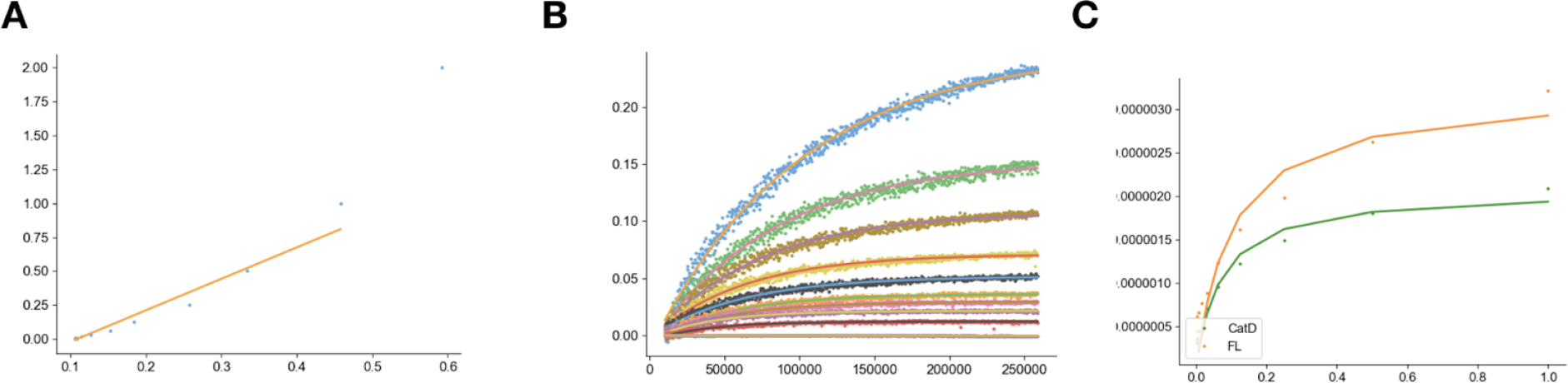
Data processing for colloidal chitin clearance assay. (a) Concentrations from enzyme-free controls were matched to absorbance, and for concentrations below 0.5% w/v a linear regression fit the data reasonably well. (b) Progress curves of a concentration series of bulk chitin were subtracted from their initial state, then fit by a non-linear relaxation analysis to extract initial rates. (c) Initial rates were plotted against substrate concentration and were fitted via non-linear regression to a Michaelis-Menten curve to extract rate constants.

**Figure S3.**
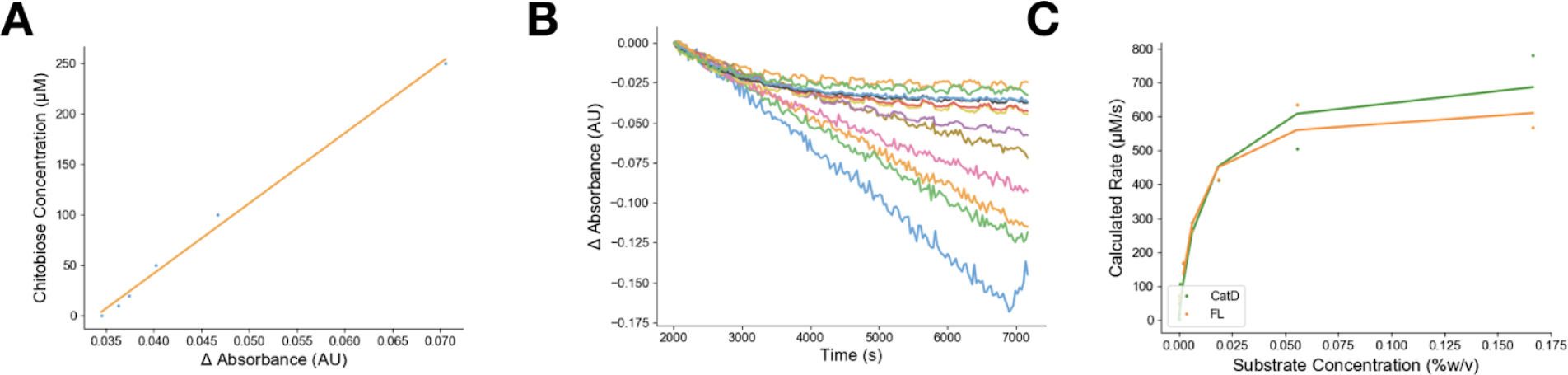
Data processing for ferricyanide reduction assay. (a) Concentrations from chitobiose controls were matched to absorbance, and for concentrations below 250 μM a linear regression was fit the data. (b) From progress curves for the non-enzymatic reaction with potassium ferricyanide, the maximum and minimum values were subtracted from each other and scaled by the incubation time to extract the rate of generation of soluble reducing sugars. (c) Rates were plotted against substrate concentration and were fitted via non-linear regression to a Michaelis-Menten curve to extract rate constants.

**Figure S4.**
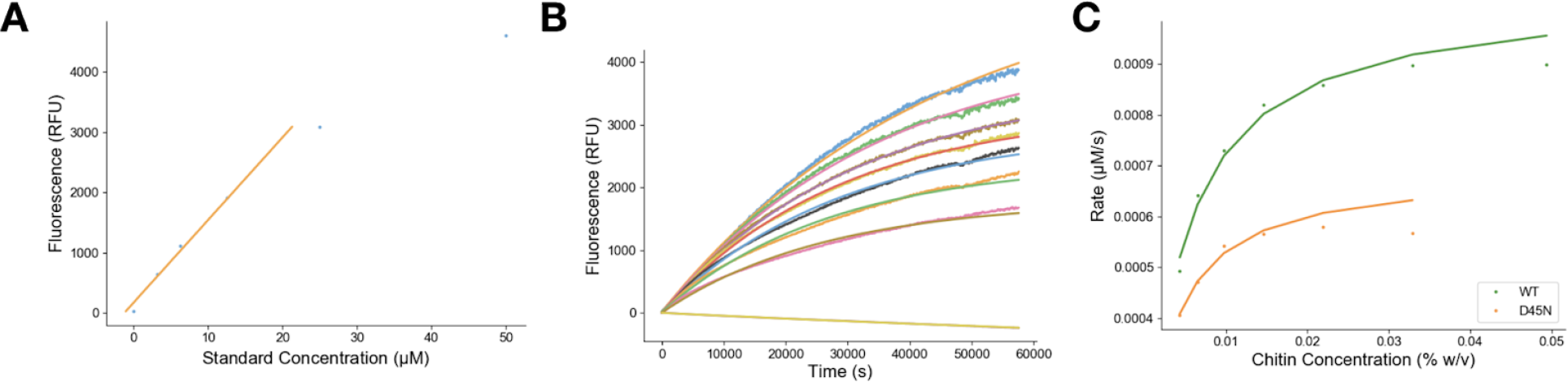
Data processing for chitO assay. (a) Concentrations from chitobiose controls were matched to fluorescence after incubation with chitO, horseradish peroxidase, and quantared, and for concentrations below 30 μM a linear regression was fit the data. (b) From progress curves, a non-linear regression was used to fit relaxation parameters to extract initial rates for a concentration series of colloidal chitin (c) Rates were plotted against substrate concentration and were fitted via non-linear regression to a Michaelis-Menten curve to extract rate constants.

## Acknowledgements

JSF was supported by a Pew Scholar Award from the Pew Charitable Trusts, a pilot grant from the Sandler Asthma Basic Research Center (SABRE Center), and NIH GM123159. BAB was supported by an ARCS scholarship, a UCSF Discovery fellowship, and by NIH GM008284. RED was supported by the National Science Foundation Graduate Research Fellowship under Grant No. (1650113). SvD and RML were supported by NIH R01 HL128903 and HHMI.

